# Defining the Impact of Cell-Type and Species on the Molecular RNA Replication Kinetics of Seoul Virus

**DOI:** 10.1101/2025.04.24.650411

**Authors:** Autumn T. LaPointe, Stefan D. Klimaj, Alison M. Kell

**Affiliations:** Department of Molecular Genetics and Microbiology, University of New Mexico School of Medicine, Albuquerque, New Mexico, United States of America

**Keywords:** Seoul Virus, RNA Replication, qRT-PCR, Negative-stranded segmented virus

## Abstract

Hantaviruses are zoonotic, tri-segmented, negative-sense RNA viruses and a significant public health threat. Viral pathogenesis varies between host species, with rodent reservoir infection being asymptomatic and human infection resulting in severe, immune-mediated disease. Viral pathogenesis is highly dependent on virus replication efficiency since it affects the virus’s ability to evade detection and determines the magnitude of the host immune response. However, the molecular replication kinetics for hantaviruses remain poorly defined. Therefore, we developed a sense- and segment-specific quantitative real-time PCR (ssqRT-PCR) assay and a SYBR-based qRT-PCR (Sb-qRT-PCR) assay, allowing us to quantify both negative-sense genome levels and total viral RNA synthesis of the small (S), medium (M), and large (L) segments of Seoul virus (SEOV). We then measured total viral RNA and genome accumulation in reservoir rat endothelial cells (RLMVEC), non-reservoir human endothelial cells (HUVEC-C), and Vero E6 epithelial cells. We also measured the ratio of each segment released into the culture supernatant, approximating the relative packaging efficiency. We found that, while the magnitude of viral RNA differed, RNA replication kinetics were largely similar between reservoir and non-reservoir endothelial cells. However, replication and release kinetics differed between infection of endothelial and Vero cells. We also found that the S, M, and L segments were not equally abundant during viral infection or release, but instead followed a trend of M>L>S. Overall, this study validates two qRT-PCR assays to measure SEOV RNA, details the accumulation and release of each viral segment, and demonstrates the impact of host cell type on hantavirus replication.

**Impact Statement:** Hantavirus infections in humans lead to significant immune-mediated morbidity and mortality around the world and pose a threat to public health. Zoonotic transmission occurs from persistently infected rodents which show no overt signs of disease. Understanding the molecular interactions that drive divergent infection outcomes is incomplete in the field of hantavirus research due to limited tools and reagents. This novel strand- and segment-specific quantitative RT-PCR assay uncovers the unique kinetics of hantavirus genome replication in reservoir and non-reservoir cells. Further, this assay will serve as a valuable tool to investigate antiviral therapies and virus-host interactions critical for polymerase activity.

## Introduction

Hantaviruses are negative-sense, segmented RNA viruses which are maintained worldwide in rodent and insectivore reservoirs [1, 2]. While infection of the rodent reservoir is typically asymptomatic and persistent, hantavirus infection of humans can result in severe disease [3]. In the case of Old World hantaviruses, disease manifests as hemorrhagic fever with renal syndrome (HFRS) with a mortality rate of up to 12% [4, 5]. Meanwhile, infection with New World hantaviruses results in hantavirus cardiopulmonary syndrome (HCPS) with a mortality rate of up to 60% [6, 7].

Hantavirus infection in both the rodent reservoir and human host primarily targets endothelial cells [8, 9]. Infection of human endothelial cells leads to immune activation in cell culture, which may contribute to overall immune-mediated disease [7, 10, 11]. For Old World hantaviruses, the RNA recognition receptors retinoic acid-inducible gene I (RIG-I) and melanoma differentiation-associated protein 5 (MDA5) are required for type I interferon (IFN) signaling [7, 11, 12]. The RNA pathogen-associated molecular pattern (PAMP) that is detected by RIG-I during hantavirus infection remains unknown.

Hantavirus infection of the rodent reservoir is asymptomatic *in vivo* and fails to drive significant type I IFN signaling after viral entry *in vitro* [1, 2, 13–15]. This lack of immune response is not due to direct antagonism of RIG-I by viral proteins [7]. It is therefore possible that the differences seen in immune response to SEOV infection between human and rat endothelial cells may be due to differential production of viral RNA PAMPs that could be detected by RIG-I. Alternatively, infection by incomplete viral particles that contain only one or two segments instead of three could also lead to immune sensing of improperly hidden PAMPs resulting in an inflammatory immune response. However, current knowledge of the RNA replication kinetics and genome release of SEOV in endothelial cells remains limited.

The SEOV genome is composed of three negative-sense RNA segments termed small (S), medium (M), and large (L) based on their respective sizes. During viral infection, two positive-sense RNA species are produced for each segment: the antigenome, which serves as template for genome replication, and the viral mRNA [16–18]. Since the genomic RNA is packaged into viral particles, it represents the product of viral replication. As such, genome abundance represents the sum of viral replication and particle release. However, since we currently do not know which viral RNA species (i.e. positive-sense or negative-sense) leads to the activation of the immune response, understanding viral RNA synthesis as a whole is also important.

In order to measure the RNA replication kinetics of SEOV, we developed a TaqMan strand-specific, quantitative, reverse-transcription, real-time PCR assay (ssqRT-PCR) to quantify viral genome abundance specifically as well as a SYBR-based quantitative, real-time PCR (Sb-qRT-PCR) assay to measure total viral RNA synthesis for each segment. Using these assays, we measured SEOV RNA accumulation and release over time in cell cultures. Rat and human endothelial cells represent the target cells for SEOV infection the reservoir and non-reservoir host, respectively. Additionally, we quantified SEOV RNA kinetics in Vero E6 cells, as they are an ubiquitous epithelial cell line used for hantavirus propagation and allowed us to measure SEOV replication in the absence of type I IFN. We hypothesized that there would be significant differences in RNA replication kinetics between the human and reservoir rat endothelial cells, but that RNA replication would be similar between the rat endothelial cells and Vero E6 cells, as neither mount a significant type I IFN response to SEOV infection. However, we instead observed viral replication in both human and rat endothelial cells to have similar kinetics to each other, yet unique compared to infected Vero E6 epithelial cells. Similar to previous reports, overall RNA abundance in reservoir endothelial cells was higher than in human endothelial cells, suggesting a general infection advantage. We also noted differential abundance of the S, M, and L segments, following a consistent pattern of M > L > S across all three cell species. Collectively, this study presents a strand-specific qRT-PCR assay used to quantify SEOV RNA replication in reservoir and non-reservoir host species and suggests that cell type may impact viral replication kinetics.

## Results

### qRT-PCR assay design

To target the negative-sense genome for cDNA synthesis, we designed cDNA primers complementary to the genome for each segment. These primers included a unique tag sequence on the 5’ end to prevent self-priming (Fig 1A) [19, 20]. Real-time PCR primers targeting the unique tag sequence (3’) and viral genome (5’) were used to amplify the cDNA and TaqMan probes specific for each segment were used for detection.

**Figure 1.**
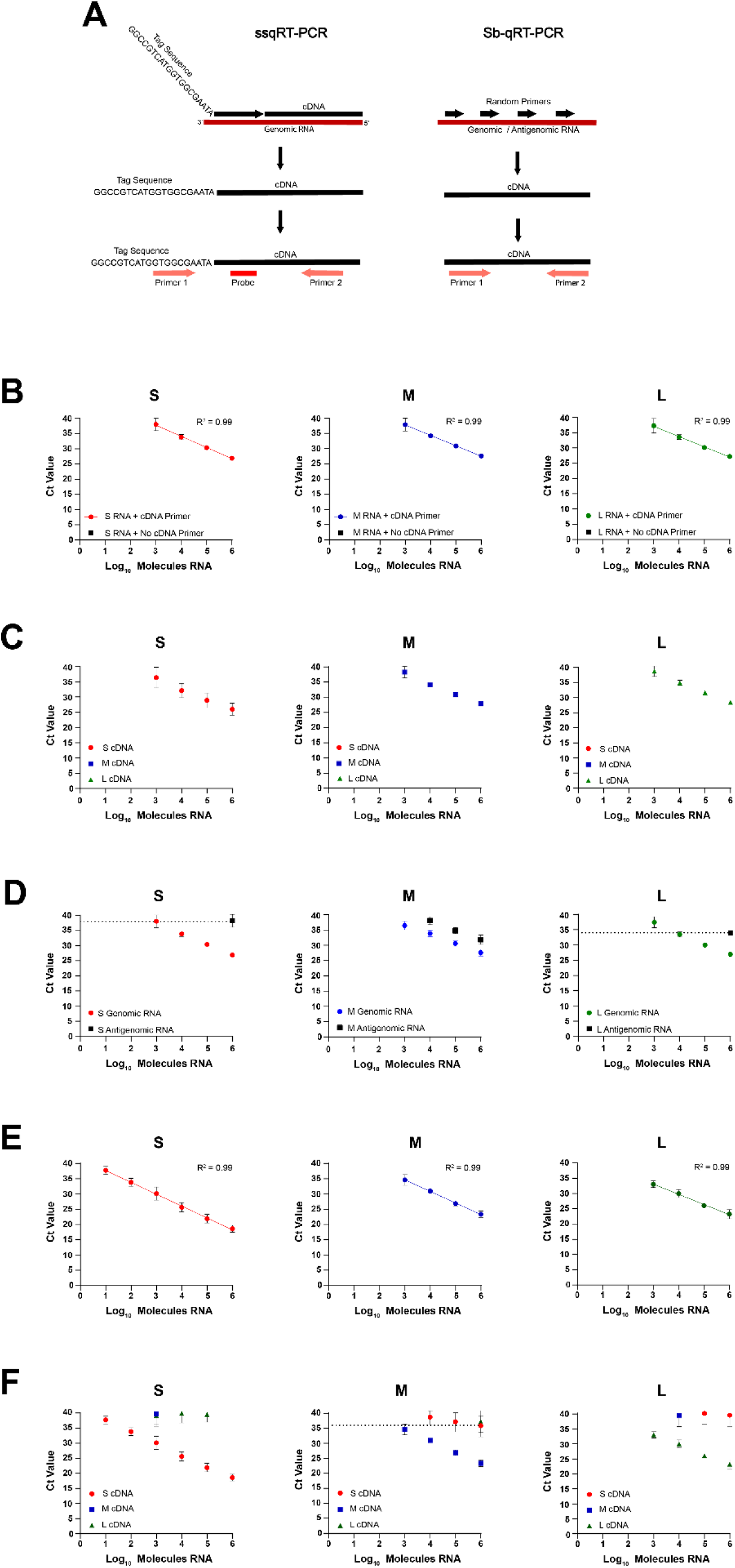
Validation of the ssqRT-PCR and Sb-qRT-PCR using *in vitro* transcribed RNAs. (A) Schematic of the cDNA synthesis and qRT-PCR strategies for the ssqRT-PCR and Sb-qRT-PCR assays. Standard curves using serial 10-fold dilutions of either S, M, or L negative-sense IVT RNAs (red circles = S, blue squares = M, green triangles = L) in the ssqRT-PCR assays (B-D) or Sb-qRT-PCR assays (E,F). (B, E) Assay sensitivity was determined by using serial dilutions of S, M, or L IVT RNA with the paired primers/probe for each assay, and dotted lines represent C_t_ cutoff for reliable template quantification using the respective assay. (C, F) Segment specificity was determined by using S, M, or L IVT RNA individually against the each segment’s primers/probe (primer/probe used is indicated in the graph titles). Data shown represent the mean of ≥3 independent experiments ±SD.

As a corollary to the ssqRT-PCR, we also designed segment-specific primers for a SYBR green assay (Sb-qRT-PCR) to measure total viral RNA (genome, antigenome, and viral mRNA) for each segment. Random primers were used for cDNA synthesis so that both the negative and positive sense RNAs are intentionally primed. Segment-specific primers were then used for the subsequent qRT-PCR step (Fig. 1A).

### qRT-PCR assay validations

To validate our assays, we *in vitro* transcribed (IVT) negative-sense RNAs for each SEOV segment from PCR templates. While we were able to transcribe the full-length S genome, we were unable to obtain full length M and L genomes, so 500nt segments spanning the primer and probe binding regions were used for the M and L genomes instead. These IVT RNAs were used as template for first-strand cDNA synthesis. Utilizing first-strand reverse transcription minimizes any bias in efficiency due to differences in template length, as these efficiency differences are not repeatedly perpetuated in a single cycle reaction as they would be in a typical, multi-cycle PCR reaction. The cDNA was then serially diluted to make a standard curve from 10^6^-10^0^ copies of input RNA for qRT-PCR. To confirm that the ssqRT-PCR assay detected only specifically primed RNA and not self-primed RNA, we included a negative control cDNA synthesis reaction without cDNA primers. For the S, M, and L negative-sense IVT RNAs, the ssqRT-PCR assay reliably detected down to 10^3^ molecules of input RNA, and when a line of best fit was applied, all three segments had R^2^ values of 0.99 indicating linear changes across the standard curve (Fig. 1B).

To test the segment specificity, IVT negative-sense RNAs of either S, M, or L were used as template for each segment’s assay (cDNA primer, qRT primers, and probe). For example, the S segment IVT RNA was used as template for the M assay primer/probe set as well as for the L assay primer/probe set (Fig 1C). For all three segments in the ssqRT-PCR assay, amplification was only detected when template RNA and primer/probe sequences were matched, demonstrating that the assays are segment specific (Fig 1C).

To validate the sense specificity of the ssqRT-PCR assay, either genomic or antigenomic IVT RNA for each segment was used as template for cDNA synthesis. For the S and L genomic primers and probes, antigenomic RNA was only detected when 10^6^ copies were added to the reaction, giving C_t_ values of 38 and 34 respectively (Fig. 1D). This correlates with an 11 C_t_ value difference in detection between the genomic and antigenomic RNAs for the S segment and a 7 C_t_ value difference in detection for the L segment. To address the potential contribution of antigenomic RNA to detection, C_t_ cutoffs were set at C_t_ 38 for S and C_t_ 34 for L, so that values falling at or below these limits would be considered background detection for future experiments. Unlike S and L, M antigenomic RNA could be detected down to 10^4^ copies with the primers intended to detect only genomic RNA and only illustrated a 4.5 C_t_ value difference in detection between the genomic and antigenomic RNAs (Fig 1D). Detection of the antigenomic M RNA remained consistent despite testing multiple different cDNA and qRT primers and PCR conditions (data not shown). Limiting detection for the M segment based on antigenome detection severely reduced the range of detection for the genome.

Therefore, the primers and probe used for the M segment are not considered to be sense-specific. This validation test was only conducted for the ssqRT-PCR, where specificity for the negative sense RNA was critical to the assay’s function, and not for the Sb-qRT-PCR assay, where both positive and negative sense RNA will be amplified and detected simultaneously.

For the Sb-qRT-PCR assay, the S segment was sensitive to 10^1^ molecules of input RNA, while the M and L segments were sensitive to 10^3^ molecules RNA (Fig. 1E). When a line of best fit was applied to the standard curves, all segments had R^2^ values of 0.99. Sb-qRT-PCR segment specificity was tested the same as the ssqRT-PCR assay, where IVT negative-sense RNAs of either S, M, or L were used as template for each segment’s assay (cDNA primer and qRT primers). For the Sb-qRT-PCR assay for S and L, non-target segments (M and L for S assay, S and M for the L assay) were detected at C_t_ values of 39 or greater, at or near the limit of detection of the assay (C_t_ 40) (Fig 1F). In addition to this, the gap in detection between the intended segment and the non-target segments was approximately 9-10C_t_ (equivalent to ∼3 log). Given that the non-target segments were measured close to the limit of detection of the assay and the large difference in C_t_ between the target and non-target segments, it was determined that detection of non-target segments was unlikely to significantly contribute to the measurement of the intended segment for S and L. The Sb-qRT-PCR assay for M detected non-target S and L segments at C_t_ values of ∼36 when 10^6 copies were added. To address the potential contribution of S and L RNA to detection of M, a limit of detection was set at C_t_ 36 for M, so that values falling at or below this limit would be considered background detection.

In order to compare SEOV replication kinetics during infection of reservoir and non-reservoir species, it was important to validate that the primers and probes designed for the ssqRT-PCR and Sb-qRT-PCR assays did not detect cellular RNAs from either the rodent reservoir or human cells. To test this, 500ng cellular RNA isolated from uninfected human umbilical vascular endothelial cells (HUVEC-C) or primary rat lung microvascular endothelial cells (RLMVEC) was used as template for cDNA synthesis followed by either ssqRT-PCR or Sb-qRT-PCR (Supp. Fig. 1A and 1B). None of the ssqRT-PCR assays detected host RNAs from either HUVEC or RLMVEC (Supp. Fig. 1A) For the M segment SB-qRT-PCR, host RNAs were detected at 39 C_t_, but these were greater than the previously set C_t_ cutoff and would be treated as background in future experiments (Supp. Fig. 1B).

We further demonstrate that the addition of non-target RNAs did not interfere with the detection of the intended viral RNAs. To do this, 150ng of either HUVEC or RLMVEC RNA was mixed with 10^6 copies of each of the genomic and antigenomic S, M, and L IVT RNAs. This mixture was then used as template for either the ssqRT-PCR or Sb-qRT-PCR assay and was compared to a standard curve for the target genome or segment. For both the ssqRT-PCR assay (Fig. 2A and 2B) and Sb-qRT-PCR (Fig. 2C and 2D), the dilution series made by the host/virus RNA mixture closely mirrored the standard curves made using a single genomic IVT RNA for all three segments, indicating that the non-target cellular or viral RNAs do not interfere with target RNA detection for either assay.

**Figure 2.**
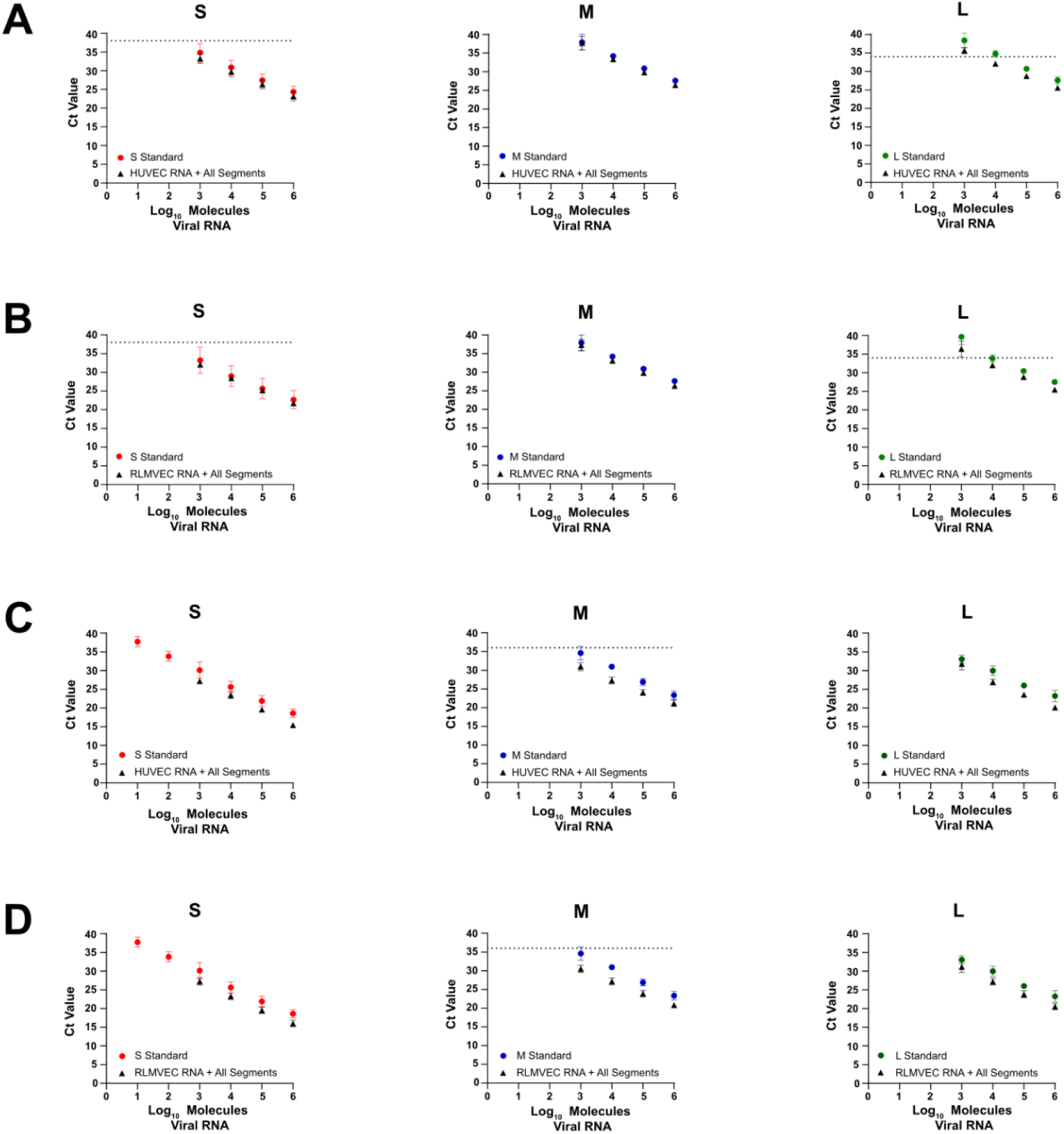
ssqRT-PCR and Sb-qPCR assays do not detect RNAs from uninfected host cells. To validate that neither the ssqRT-PCR assay (A, B) nor the Sb-qRT-PCR assay (C,D) detected host cellular RNAs, standard curves were made using serial dilutions of S, M, or L negative-sense IVT RNA (circles) or a mixture of 10^6^ copies per strand of S, M, and L genomic and antigenomic IVT RNAs plus 150ng of either uninfected HUVEC (A,C) or uninfected RLMVEC (B, D) RNA (triangles). The IVT RNA alone or mixture of IVT RNAs and uninfected cellular RNA was then used as input for either the ssqRT-PCR assay (A,B) or SB-qRT-PCR (C,D) assay using the primers/probe indicated in the graph title. Data shown represent the mean of ≥3 independent experiments ±SD.

### Quantification of viral RNA in SEOV infection

To measure SEOV RNA replication kinetics, HUVEC, RLMVEC, and Vero E6 cells were infected with a cell-specific multiplicity of infection (MOI) of 0.1 focus-forming units (FFU)/cell. Cell-specific MOI was determined as previously described to ensure that each cell type received the infectious units required for 10% infection of the culture [7]. This was done to account for differences in cell susceptibility to infection, allowing us to study RNA accumulation from a more equivalent baseline of infection. Cells were harvested every 24hrs over three days to capture viral RNA accumulation. Because the ssqRT-PCR assay to detect M genome was not sense-specific, only the S and L genomes were quantified in SEOV-infected cells. Viral replication in HUVEC displayed a ∼3.7 fold increase in S genomic RNA from 0dpi to 1dpi and a ∼6.7 fold increase in S genomic RNA, while the L genomic RNA increased ∼1.7 fold from 0dpi to 1dpi and increased ∼2.8 fold from 1dpi to 2dpi (Fig. 3A, Supp. Table 1). After 2dpi, both S and L genomic abundance in HUVEC plateaued. S and L genomic RNA levels in infected RLMVEC increased ∼101 fold for S and ∼65 fold for L from 0dpi to 1dpi, but then remained largely the same for the remainder of the infection. Conversely, infection of Vero cells resulted in increases in S and L genomic RNA across all three days. The S genome levels in infected Vero cells increased ∼9 fold from 0dpi to 1dpi, ∼5 fold from 1dpi to 2dpi, and ∼3.5 fold from 2dpi to 3dpi. Meanwhile L genome levels increased ∼5 fold from 0dpi to 1dpi, ∼6 fold from 1dpi to 2dpi, and ∼3.5 fold from 2dpi to 3dpi.

**Figure 3.**
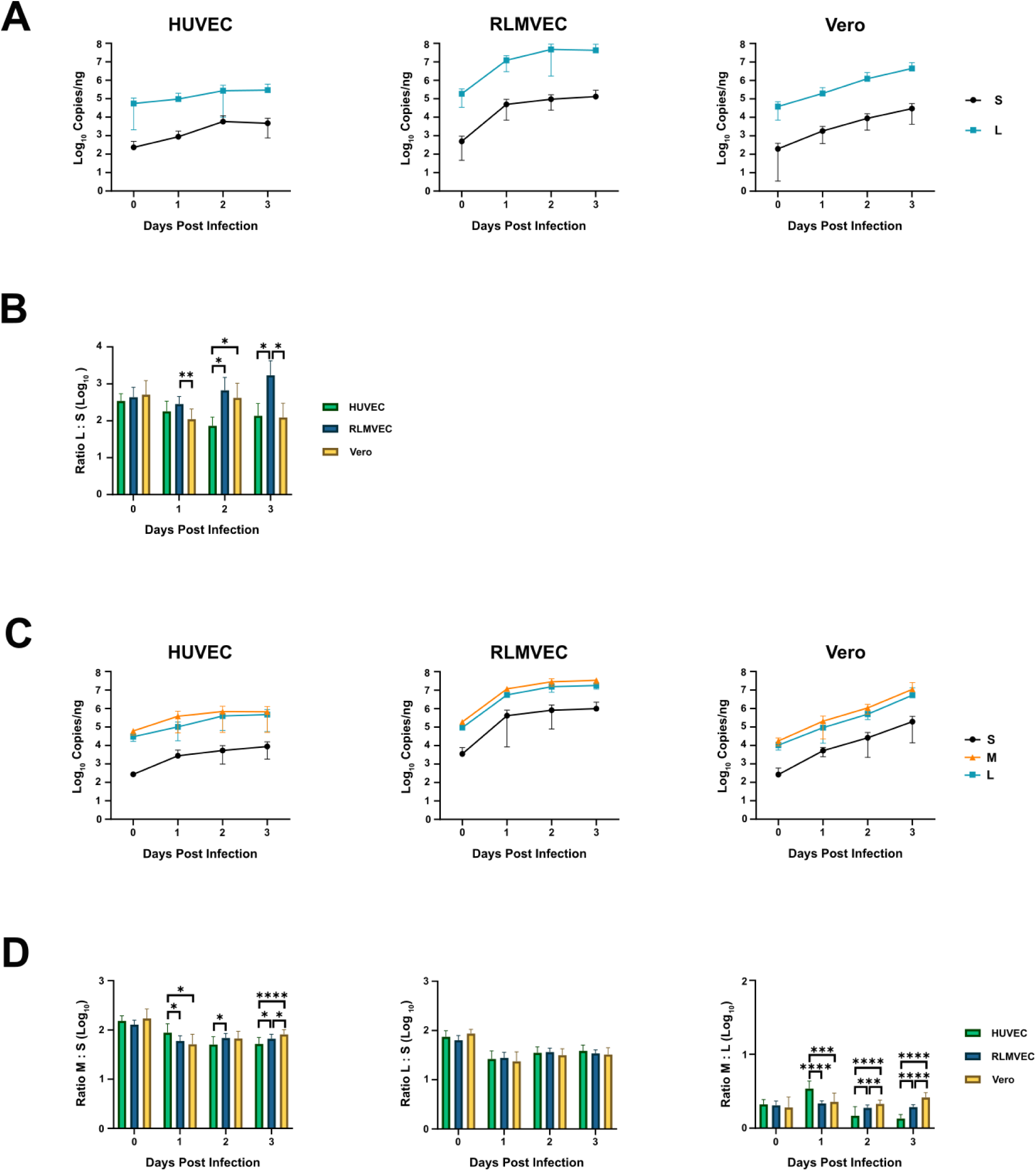
Quantification of viral RNA in SEOV-infected reservoir and non-reservoir cells. HUVEC, RLMVEC, and Vero cells were infected with SEOV at an MOI of 0.1 FFU/cell. Cells were harvested and RNA extracted at the indicated days post infection. (A) S and L genomic RNA accumulation was measured using ssqRT-PCR for each cell type. (B) The ratio of L RNA copies to S RNA copies was then calculated for each time point. (C) Combined positive and negative sense viral RNA accumulation was measured using Sb-qRT-PCR for all three segments. (D) The ratios of M RNA copies to S RNA copies, L RNA copies to S RNA copies, and M RNA copies to L RNA copies were then calculated for each time point. Data shown represent the mean of ≥3 independent experiments ±SD. Statistical significance determined by Student’s t test. *, p<0.05; **, p<0.01; ***, p<0.001, ****, p<0.0001.

**Table 1.**
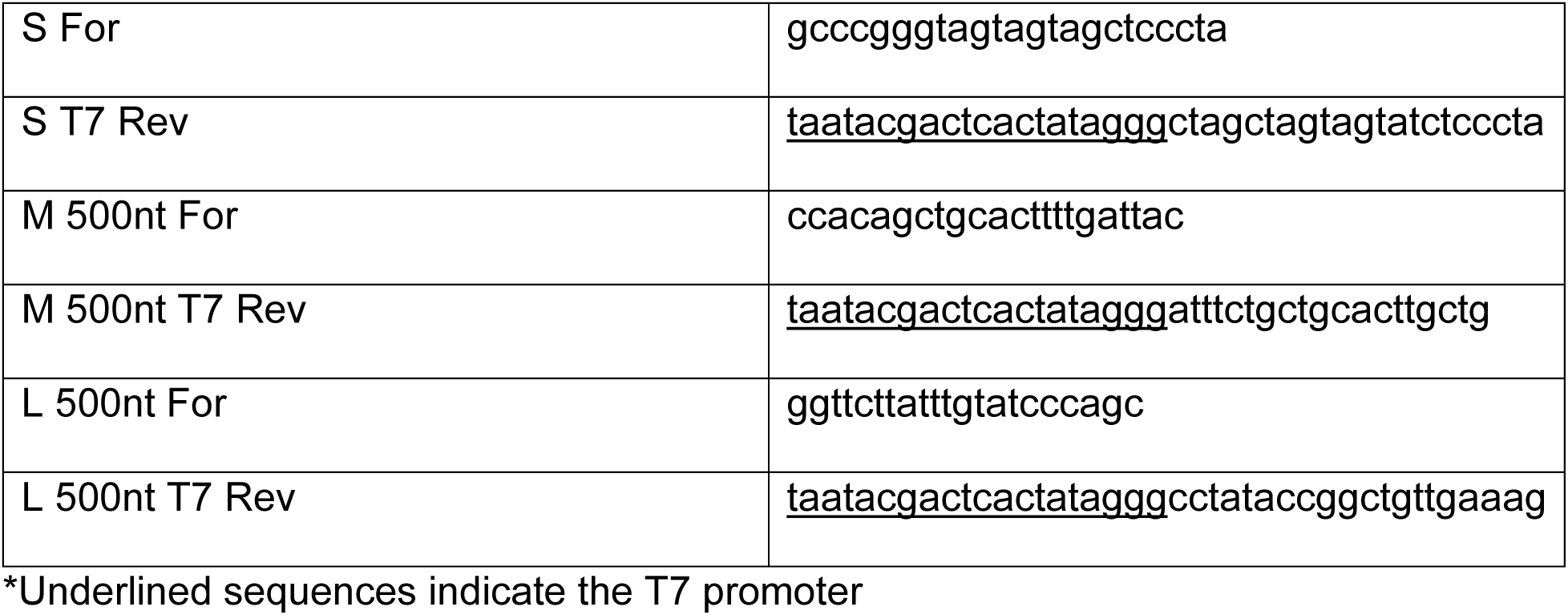
PCR Primers for Genomic IVT templates.

Cumulatively, from 1dpi to 3dpi, the S genome levels increased ∼16.4 fold and L genome increased ∼22.4 fold in infected Vero cells.

The ratio of L genome to S genome produced was also calculated for each cell type over the course of infection. For all three cell cultures, the L genome was found to be more abundant than the S genome at all timepoints. In general, the significantly higher ratio of L : S produced during infection of RLMVEC means a greater abundance of L genome was produced relative to S genome compared to the HUVEC or Vero cells (Fig. 3B, Supp. Table 2). Additionally, the ratio of L : S in RLMVEC increased over the course of infection, while the ratio of L : S remained more equivalent across the three days for infected HUVEC and Vero cells.

**Table 2.**
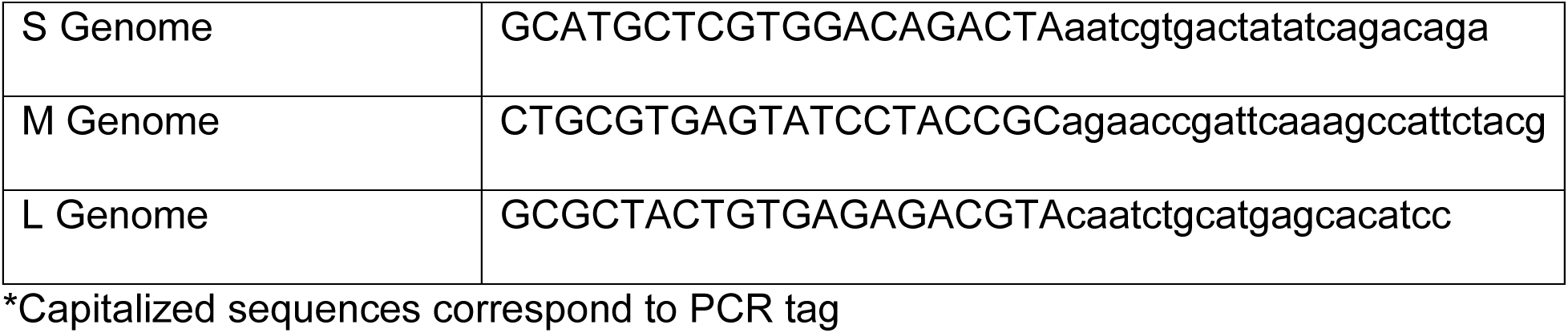
cDNA Primers.

We also used the Sb-qRT-PCR assay to measure the combined accumulation of the genome, antigenome, and mRNA for each segment. This allowed us to quantify viral RNA synthesis as a whole as opposed to viral replication of the genome alone. Again, we observed that, after an initial increase in abundance between 0dpi and 1dpi, the abundance for all three segments remained steady in HUVEC and RLMVEC (Fig. 3C, Supp. Table 3). S, M, and L RNA levels increased ∼12 fold, ∼6.3 fold, and ∼3.4 fold form respectively in infected HUVEC, but then only increased ∼2.4 fold, ∼1.7 fold, and ∼4.6 fold respectively between 1dpi and 3dpi in infected HUVEC. Meanwhile in infected RLMVEC, S, M, and L RNA levels increased ∼117 fold, ∼61.6 fold, and ∼58.6 fold respectively from 0dpi to 1dpi, but then increased only ∼2.1 fold, ∼3 fold, and ∼3.1 fold respectively from 1dpi to 3dpi. Consistent increases in abundance for all three segments were observed during Vero cell infection. S, M and L RNA levels increased ∼19.6, ∼11.7, and ∼8.8 fold respectively from 0dpi to 1dpi. All three segments increased ∼5 fold from 1dpi to 2dpi and both M and L RNA levels increased ∼10 fold from 2dpi to 3dpi, while S RNA levels increased ∼7.5 fold from 2dpi to 3dpi. Cumulatively, from 1dpi to 3dpi RNA levels increased ∼40 fold, ∼50 fold, and ∼56 fold for S, M, and L respectively in the infected Veros. The M segment was the most abundant in all three host species, followed by L segment and then the S segment. Calculating the ratio of M : S for each cell culture found largely similar relative abundances (Fig. 3D, Supp. Table 4). Likewise, the ratio of L : S and M : L were also remarkably similar between cell cultures and over the course of infection. Although some of these ratios were found to be statistically significantly different between cell types, the magnitude of the difference is modest. Overall, while the RNA accumulation for each segment suggests that SEOV RNA replication may differ between these cells, the ratios between segments remain largely unchanged over the course of infection and between cell types.

**Table 3.**
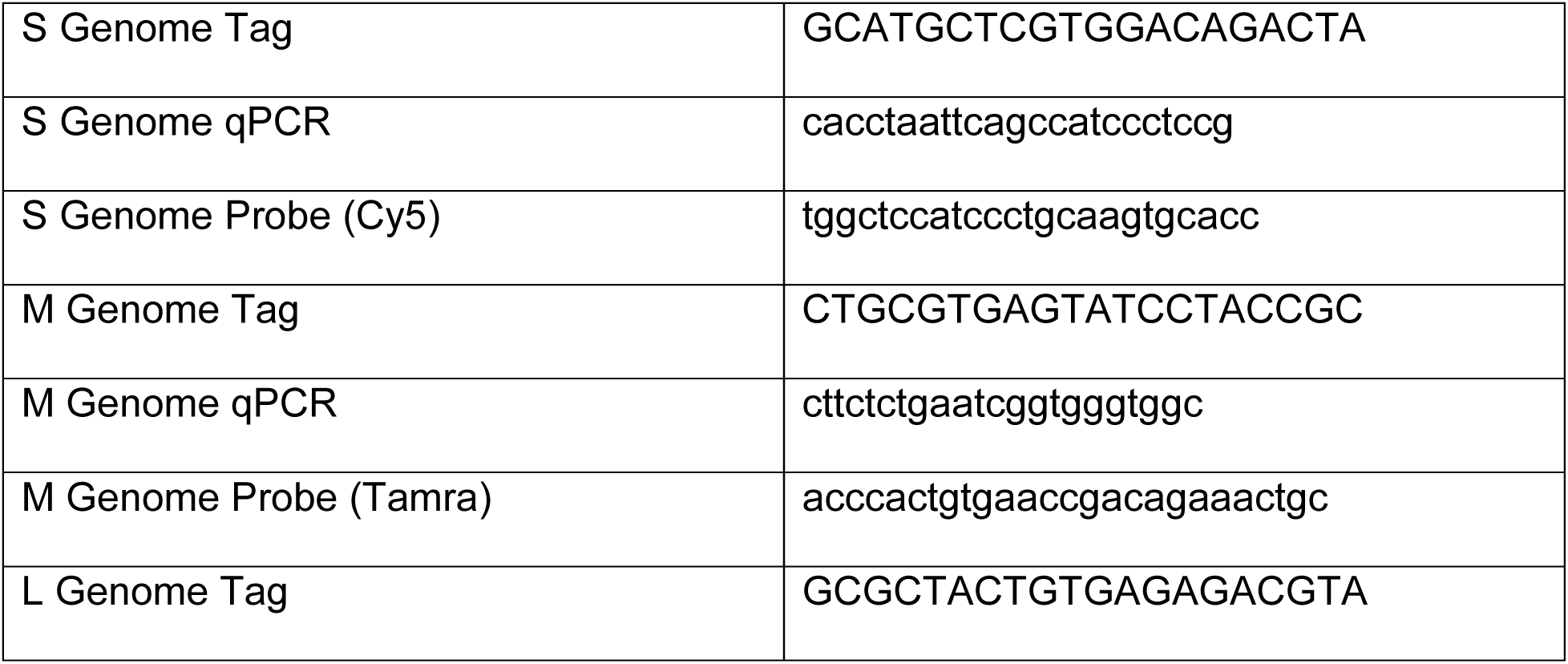
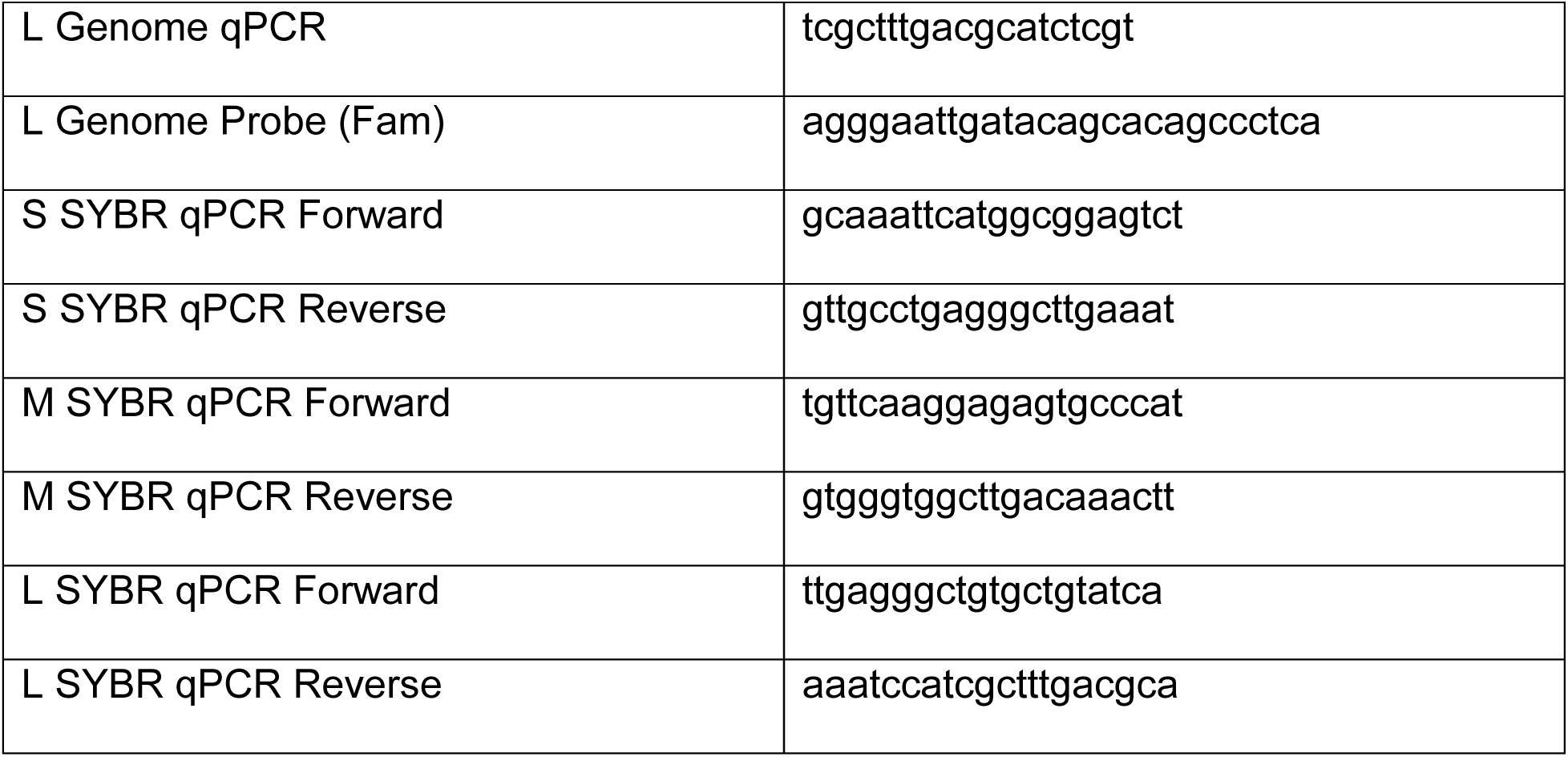
qRT-PCR Primers and Probe.

To determine whether the trends observed in genomic and total SEOV RNA synthesis were also observed in particle production, supernatants from the above infected cells were collected. A focus-forming unit (FFU) assay was used to determine viral titer while the ssqRT-PCR assay was used to measure genome release. Trends in viral titer largely follow what was observed for viral RNA replication, with titers from the human and rat endothelial cells increasing 2.5 and 4.5 fold respectively from 1dpi to 2dpi and then plateauing (Fig. 4A). Titer from Vero cells increased ∼10.6 fold from 1dpi to 2dpi and then ∼2.3 fold from 2dpi to 3dpi. To measure genome copies released from infected cells, viral particles were purified by ultracentrifugation through a 30% sucrose cushion to remove cell debris and naked RNA that might influence the ssqRT-PCR assay. Although the ssqRT-PCR was not used for measuring intracellular M genome in Figure 3, this assay was used to measure the amount of M genome being released, as the negative-sense genome is the dominant RNA species in bunyavirus particles [21]. Importantly, this assay does not determine which segments are packaged together in a single virion, but simply the rate at which each segment is released in particles from the cell. The quantified abundance of each segment released largely reflected our intracellular observations, with M being the most abundant, followed by L, then S (Fig. 4B). S segment abundance increased during infection of HUVEC between days 1 and 2 post infection ∼25 fold and then plateaued, while M segment release remained equivalent between 1dpi and 2dpi and then increased ∼6.5 fold form 2dpi to 3dpi (Supp. Table 5). L segment abundance remained steady across all three days. For infected RLMVEC, S segment abundance increased ∼10 fold from 1dpi to 2dpi, then decreased ∼2.5 fold from 2dpi to 3dpi. Meanwhile the M segment remained steady over all three days and the L segment only increased ∼2.8 fold between 1dpi to 2dpi. S genome release from infected Vero cells increased consistently across all three days, ∼4 fold from 1dpi to 2dpi and ∼6.3 fold from 2dpi to 3dpi. M genome release increased ∼2.6 fold from 2dpi to 3dpi and L genome increased ∼2.8 fold from 1dpi to 2dpi then remained steady. Calculating the ratio of each segment released, the ratio of M : L segment released from HUVEC was significantly smaller compared to the ratio of M : L released from RLMVEC and Vero cells on days 1 and 2 post infection (Fig. 4C, Supp. Table 6).

**Figure 4.**
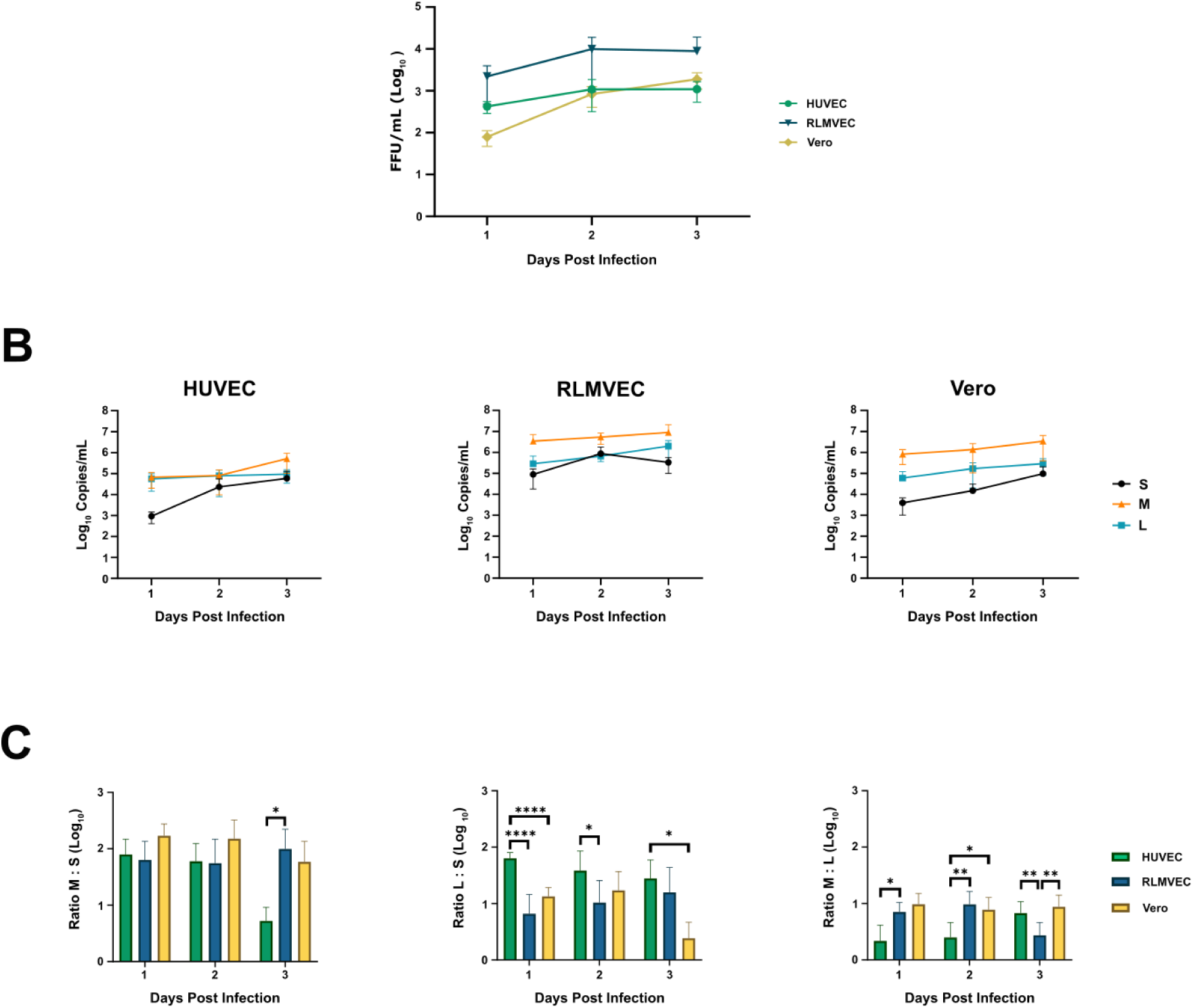
Quantification of infectious virus and RNA genome release from SEOV-infected cells. HUVEC, RLMVEC, and Vero cells were infected with SEOV at an MOI of 0.1 FFU/cell. (A, B) Supernatants were collected at the indicated times post infection and used for either measuring viral titer via FFU assay (A) or RNA extracted to measure genomic RNA via the ssqRT-PCR assay (B). (C) Ratios of M RNA copies to S RNA copies, L RNA copies to S RNA copies, and M RNA copies to L RNA copies were calculated for each time point. Data shown represent the mean of ≥3 independent experiments ±SD. Statistical significance determined by Student’s t test. *, p<0.05; **, p<0.01; ***, p<0.001, ****, p<0.0001.

Infected HUVEC also released a significantly larger ratio of L : S. Meanwhile, the ratios of M : S, L : S, and M : L released from infected RLMVEC and Vero cells are similar between the two cell types, with the exception of M : L at 3dpi where the ratio was significantly smaller for RLMVEC infection compared to Vero.

## Discussion

The strand-specific qRT-PCR assay can specifically measure the negative sense genome segments of SEOV, and represents the first system to do so for a hantavirus.

To complement this assay and obtain a holistic view of SEOV replication, we also designed a SYBR based qRT-PCR assay that measures the total viral RNA produced for each segment. The ssqRT-PCR and Sb-qRT-PCR assays allowed us to conduct an analysis of SEOV RNA replication and release in multiple cell types. This work builds on previous studies in the hantavirus field which used qRT-PCR to characterize the production of viral mRNA in Sin Nombre virus (SNV) and total viral RNA synthesis and release in Puumala virus (PUUV) [18, 22]. While our study of SEOV total RNA replication indicated that M segment was the most abundant in cells, followed by L and then S, SNV and PUUV were reported to produce the S segment most abundantly, followed by M and then L. The MOI used in these studies could be a technical explanation for this discrepancy. The previous studies used an MOI of 6 or 10 infectious units per cell in order to quantify viral RNA replication within the first hours of infection. Conversely, we conducted infections at an MOI of 0.1 FFU/cell to limit the potential impact of defective interfering particles. Using a higher MOI, while potentially necessary to measure RNA replication at early timepoints, could lead to multiple simultaneous entry events and replication of defective genomes [23–25]. This could result in increased abundance of a particular segment that would not occur if only one infectious unit entered a cell. An additional biological explanation for the differences we see in segment abundances is that SNV and PUUV genomes encode an additional nonstructural protein on their S segment, termed NSs, not found in SEOV [26, 27].

While NSs expression or putative sequences have been observed for hantaviruses with rodent reservoirs in the Cricetidae family (e.g. SNV, Andes virus, and PUUV), hantaviruses that infect rodents from the Muridae family (e.g. SEOV and Hantaan virus) lack an NSs [28, 29]. Since both SNV and PUUV encode both the NSs and N on the S segment, it is possible that hantaviruses with an NSs might produce more S segment than those which only encode one protein on the S segment. A more extensive study measuring the RNA replication of multiple hantaviruses at different MOIs and times post infection is necessary to clarify the relative differences seen in segment abundance between these studies. Interestingly, a recent study involving distantly related phleboviruses, found the M genome untranslated regions (UTR) of Heartland virus to be the most efficient for transcription and replication, followed by the L UTR and then the S UTR [30]. Future work applying the SEOV minigenome system could determine whether this is also true for hantaviruses and might explain our findings here [31]. Furthermore, while the current study focused on utilizing the ssqRT-PCR and Sb-qRT-PCR assays to measure SEOV, they have not been validated as specific for SEOV compared to other hantaviruses and therefore should not be used in a diagnostic setting. Determining the viral specificity of these assays remains an avenue for future work.

Notably, using the SYBR assay, the ratio of total RNA species quantified between segments remained fairly similar over the course of infection, especially the ratio of L : S. This suggests that total RNA of each segment is produced at similar rates. The fact that the ratios of the total RNA produced for each segment are largely consistent during infection means that the differences seen in RNA copy abundance between segments are not due to the rates at which each segment is produced, but instead are likely due to early infection events. One potential explanation could be unequal packaging of S, M, and L genomes into particles in the viral stocks, which could lead to differences in RNA abundance after viral entry. However, for this to be the sole cause for the differences in abundance between each segment, there would need to be an almost two log difference between incoming genome segments. Therefore, unequal packaging of genomic segments in the viral stock particles is unlikely to be the sole contributing factor. Alternatively, it is also possible that replication of the S, M, and L segments is initiated at different times, leading to the difference seen in RNA abundance. This idea that replication of each segment may start at different times is supported by Hutchinson et al, who observed that amplification of the S mRNA was the first to be detected at 4hpi, followed by M mRNA at 8hpi, and then L mRNA at 48hpi in SNV infection [18]. Minor differences in segment ratios for total viral RNA synthesis over the course of infection of all three cell types suggests a conserved mechanism for regulating viral RNA production. Therefore, targeting these conserved mechanisms used for RNA replication or the pathways that dictate when replication of each segment is initiated may be useful strategies for antiviral development.

Similar to what has been observed for PUUV, general trends in genome release reflect total intracellular viral RNA [22]. We observed that in both genome release and intracellular RNA synthesis M is the most abundant followed by L and S. We also observed similar kinetic trends, with infected HUVEC and RLMVEC showing slight increases in genome release, but consistently larger increases in genome release as Vero infection progressed. However, one interesting difference observed was that the abundance of M and L genomes that were released from infected HUVEC are very similar compared to infection of the other two cell types, where M and L genome abundance differ by a log or more. Future studies will be needed to determine if these differences in SEOV genome release are solely immune-mediated or result from differences in host : virus interactions.

While it was unexpected to find that the segments were released in different ratios, it is important to note that the qRT-PCR assays used do not determine co-packaging or whether the RNA is replication competent, but only the abundance of each segment in the purified supernatant. It is possible that the abundance of replication competent / infectious RNA for each segment are similar, and that the increased abundance of M and L RNA in the supernatant are representative of increased defective RNA production for those segments. It is also possible that additional copies of M and L are needed to efficiently initiate viral replication in a newly infected cell. In a previous study by Bermudez-Mendez et al using the orthobunyavirus, Schmallenberg Virus (SBV), the authors found relative segment abundance to be dependent on the cell line infected [32]. Specifically, Vero cells infected with SBV were found to release different abundances of the S, M, and L genomes, but infection of C6/36 mosquito cells resulted in the release of more equivalent amounts of all three segments. Therefore, it is possible that relative segment abundance may be context- and virus-dependent, and warrants future study.

A further consideration is the possibility that purified supernatants contain viral particles along with extracellular vesicles. Endothelial cells have been reported to generate extracellular vesicles for communication with other cells and viral RNA has been found within extracellular vesicles in other viral systems. Thus, the composition of particles in the cellular supernatant may influence the measured abundance of each segment and may be found to play a role in infection dynamics and pathogenesis for hantaviruses in the future.

Contrary to our hypothesis, although SEOV RNA was more abundant in the rat endothelial cells compared to human endothelial cells, a finding consistent with previous literature, the temporal kinetics of RNA accumulation and release remained similar between the two species [33]. The human and rat endothelial cells reach maximum RNA accumulation and viral titer by 2dpi. In contrast, the Vero cells show consistent increases throughout infection. These data suggest that SEOV may replicate differently in endothelial cells versus epithelial cells. However, it is important to note that infection of Vero cells, while epithelial, are not representative of what infection of reservoir or non-reservoir epithelial cells, as Vero cells lack a type I IFN response. Therefore, analysis in additional relevant epithelial cell types and additional strains of SEOV would be of interest. The lack of a strong type I IFN response during SEOV infection of RLMVEC and Vero cells suggests that these temporal differences are not driven solely by innate immune responses, but rather may be cell type specific [7]. Additionally, SEOV has been proposed to use the same receptors, αVβ3 integrins, on both Vero and endothelial cells, indicating that receptor usage may not explain this difference [34].

Therefore, it is possible that the variance seen in replication kinetics between the endothelial cells and the epithelial cells are due to the unique virus : host interactions that occur within each cell type. As such, future work will focus on defining virus : host interactions unique to reservoir and non-reservoir infections and their consequences for SEOV replication.

## Materials and Methods

### *In Vitro* Transcription of SEOV RNAs

Full length SEOV S genomic RNA was *in vitro* transcribed from PCR templates using the T7 MEGAscript kit (Ambion AM1334) according to manufacturer’s instructions. Full length M and L genomes were unable to be obtained with this method, so 500nt segments spanning the primer and probe binding regions were PCR amplified using the primers in table 1 (underlined portions indicate T7 promoter). These PCR templates were then *in vitro* transcribed using the T7 MEGAshortscript kit (Ambion AM1354). For full length antigenomic SEOV RNA, plasmids containing each of the positive sense RNA segments with an upstream T7 promoter sequence were linearized with NheI and then used as templates for *in vitro* transcription using the T7 MEGAscript kit according to manufacturer’s instructions.

To assess RNA integrity, RNA ladder (Thermo Scientific High Range RiboRuler FERSM1821) or IVT RNA was mixed with RNA loading dye containing ethidium bromide, heated to 80°C for 10 minutes, placed immediately on ice, and then run at 30V on a 2% formaldehyde-agarose gel. RNA bands were visualized using a BioRad ChemiDoc imaging system. Concentrations of the RNA transcripts were measured using spectrophotometry and the molecules RNA/uL for each transcript was calculated using the total molecular weight of each RNA segment.

### cDNA Synthesis

IVT RNA was mixed with 50uM dNTPs and either 1uM (S and M) or 0.1uM (L) cDNA primer (Table 2, uppercase indicates tag sequence), heated at 95C for 5min, and then immediately moved to ice. RT buffer (Maxima H-minus reverse transcription kit, Thermo Scientific K1652) and RNAse inhibitor (Abmgood G138) were then added to the reaction. The reaction was then moved to either 60C (M and L) or 65C (S) and Maxima H-minus reverse transcriptase was added. Reactions were then moved to 65C for 30min, followed by 85C for 5min, then cooled to 4C. RNase H (NEB M0297L) was added to each reaction and incubated at 37C for 20min. To measure cDNA synthesis resulting from self-priming RNAs, cDNA synthesis was carried out in the absence of the cDNA primer. For both the validation assays using host RNAs and to experimentally measure intracellular viral RNA, 500ng cellular RNA were used as input. To measure genomic RNA release, 25% of RNA extracted from pelleted supernatants was used as input.

cDNA synthesis for total viral RNA SYBR qRT-PCR was done using a high capacity cDNA reverse transcription kit (Applied Biosystems 4368814) following manufacturer’s instructions. For both the validation assays using host RNAs and to measure intracellular viral RNA, 500ng cellular RNA were used as input. Validation assays using mixed host and IVT RNAs used 150ng cellular RNA as input.

### qRT-PCR

The ssqRT-PCR assay was performed using 300nM segment-specific qPCR primer, 300nM segment specific tag primer, 200nM Taqman probe, and PrimeTime Gene Expression Master Mix (IDT 1055771; Table 3). Standard curves were generated using cDNA made from IVT RNAs such that a range of 1x10^6 copies to 1 copy cDNA were added to the ssqRT-PCR reaction. Thermocycling conditions for S: 95C for 3min, 40 cycles of 95C for 15sec and 70C for 1min. Thermocycling conditions for M and L: 95C for 3min, 40 cycles of 95C for 15sec and 60C for 1min. For infections, copy number/ng and copy number/mL was calculated using a standard curve from IVT genomic RNA. SYBR qRT-PCR was performed using Applied Biosystems SYBR Green Universal master mix (4309155) following manufacturer’s guidelines. cDNA was diluted to within range of the standard curve then mixed with 12.5nM segment specific forward and reverse primers and 1x SYBR mix. Thermocycling conditions: 95C for 10min, 40 cycles of 95C for 15sec and 60C for 1min. For infections, copy number/ng and copy number/mL was calculated using a standard curve from IVT genomic RNA.

### Cell Culture and Virus Propagation

Vero E6 cells (ATCC, CRL-1586) were cultured in Dulbecco’s modified Eagle’s medium (DMEM) supplemented with 1% pen/strep, 1% nonessential amino acids, 10% heat inactivated FBS, and 2.5% HEPES. Primary rat lung microvascular endothelial cells (RLMVEC, VEC Technologies) were cultured in MCDB-131 base medium (Corning) supplemented with VascuLife VEGF LifeFactors (Lifeline Cell Technology LS-1020) and 10% heat inactivated FBS. Human umbilical vein endothelial cells (HUVEC-C; ATCC, CRL-1730) were cultured in VascuLife EnGS Endothelial medium (Lifeline Cell Technology LM-0002) with VascuLife EnGS LifeFactors (Lifeline Cell Technology LS-1019) and 10% heat inactivated FBS. All cells were cultured at 37C and 5% CO_2_. Endothelial cells were cultured on tissue culture treated plates coated with rat tail collagen (VWR, 47747-218).

To propagate virus, Vero E6 cells were infected with Seoul virus (SEOV) strain SR11 for 12 days (MOI 0.01). Supernatant was then collected and clarified by centrifugation at 1000xg for 10min.

### Virus Infections

RLMVEC, HUVEC, and Vero E6 cells were infected with SEOV at a cell-specific multiplicity of infection (MOI) of 0.1 focus forming units / cell. Cell specific MOI was determined previously [7]. After a 1hr adsorption period at 37C, the inoculum was removed, and the cells were washed twice with 1x phosphate buffered saline (PBS) to remove unbound virus. Each cell line was given its respective media and incubated at 37C and 5% CO_2_. At the indicated times post infection, supernatant was collected and clarified at 1000xg for 10min. An aliquot (25%, 500uL) of supernatant was kept for measuring viral titer while the rest (75%, 1.5mL) was inactivated by applying UV radiation at 104,000 uJ/cm^2^ (FisherBrand UV crosslinker, 13-245-221). Virus from UV inactivated supernatant was pelleted though a 30% w/v sucrose cushion in TNE buffer (10mM Tris-HCl pH 7.5, 100mM NaCl, 1mM EDTA) for 2hrs at 30,000rpm. TRIzol reagent was added to the pellets and total RNA was extracted via isolation following manufacturer instructions. Cell monolayers were washed with 1x PBS and harvested in TRIzol reagent (ThermoFisher Scientific, 15596026). Total RNA was extracted via isolation following manufacturer’s instructions.

### Focus Forming Unit Assay

Standard virological focus forming unit (FFU) assays were used to determine the infectious titer of all viral samples as previously described [11]. Briefly, Vero E6 cells were inoculated with 5-fold or 10-fold serial dilutions of virus containing samples. After a 1hr adsorption period, cells were overlaid with 2% methylcellulose supplemented with 2x DMEM, 2% FBS, 1% pen/strep, and 1% HEPES. At 7dpi, cells were fixed with 95% EtOH : 5% acetic acid and probed with primary anti-SEOV N (custom, Genscript) and HRP-conjugated goat anti-mouse antibodies (JacksonImmunoresearch, 115-035-003). Foci were visualized using Vector VIP peroxidase substrate kit (Vector Laboratories, SK-4600).

## Author Contributions

Conceptualization: A.T.L and A.M.K; investigation: A.T.L., S.D.K., and A.M.K.; writing original draft: A.T.L. and A.M.K.; review and editing: A.T.L., S.D.K., and A.M.K.; supervision: A.M.K.

## Conflicts of Interest

The authors declare that there are no conflicts of interest.

## Funding Information

This work was supported by the National Institutes of Health NIAID grant 1R01AI171289-01 granted to AMK. ATL and SDK were supported by fellowships by the National Institutes of Health NIGMS T32AI007538. The funders had no role in study design, data collection and analysis, decision to publish, or preparation of the manuscript.

## Acknowledgements

We would like to acknowledge Dr. Dave Peabody for the use of space and helpful feedback.

---

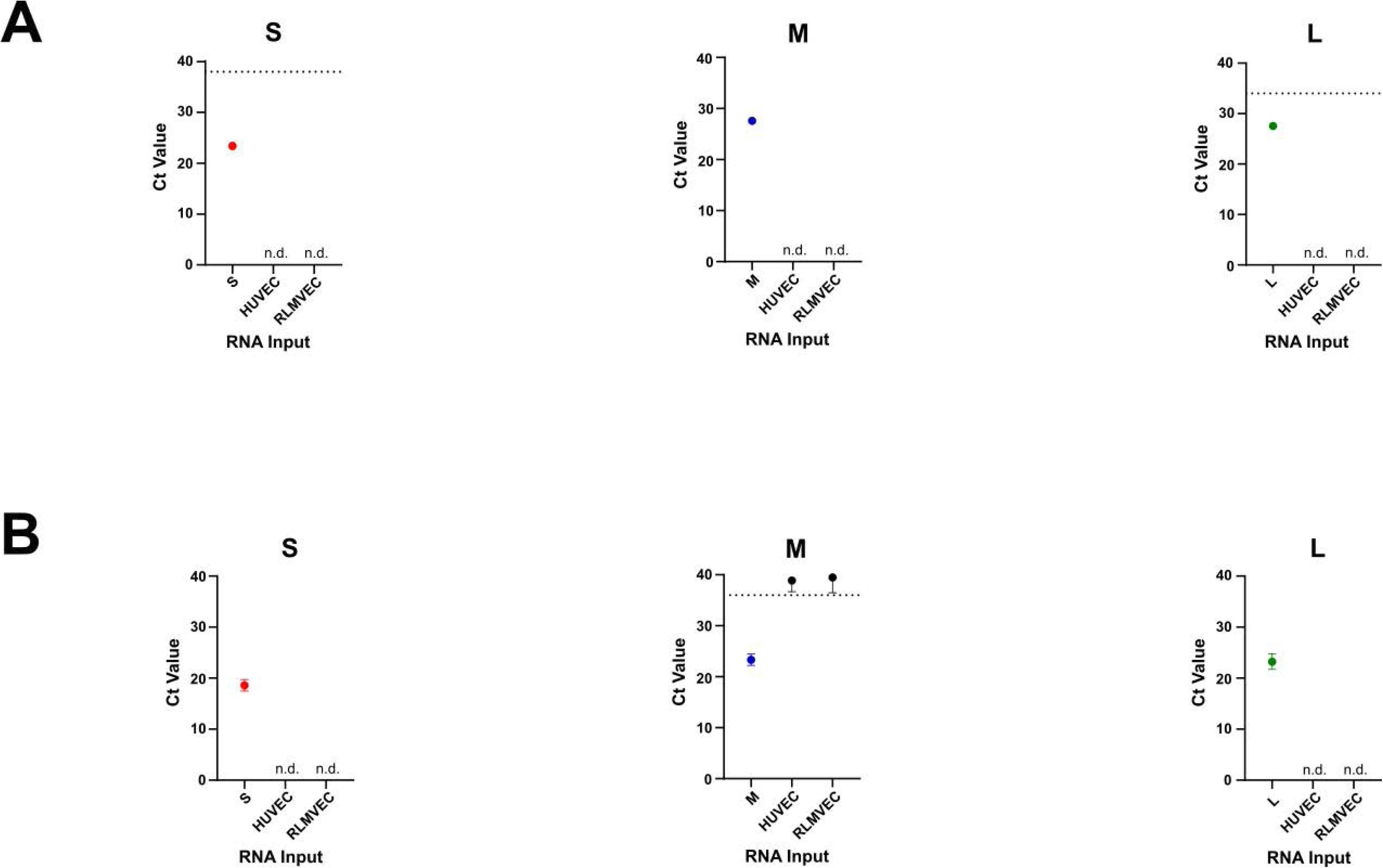

## Notes

### Competing Interest Statement

The authors have declared no competing interest.

### Summary of Updates

Author funding updated, supplementary data updated

